# Neuroanatomical Correlates in Bilinguals: The Case of Children and Elderly

**DOI:** 10.1101/586768

**Authors:** Lorna García-Pentón, Yuriem Fernández, Jon Andoni Duñabeitia, Alejandro Pérez, Manuel Carreiras

**Affiliations:** University of Toronto, Scarborough campus (UTSC), 1265 Military Trail, Scarborough, ON M1C1A4, Canada; Basque Center on Cognition, Brain and Language (BCBL), Mikeletegi 69, 2°, 20009 Donostia-San Sebastián, Gipuzkoa, Spain; University of the Basque Country UPV/EHU, Barrio Sarriena s/n, 48940 Leioa, Bizkaia, Spain; Universidad Nebrija, Santa Cruz de Marcenado, 27, 28015, Madrid, Spain; IKERBASQUE. Basque Foundation for Science, Alameda Urquijo, 36-5, 48011 Bilbao, Bizkaia, Spain

## Abstract

How bilingualism modulates brain areas beyond the language regions is still controversial. Through a comprehensive set of analyses on brain structure, we investigated brain differences between Basque-Spanish bilinguals and monolinguals in children and the elderly, the most sensitive target groups to detect potential brain differences. In particular, we employed Diffusion MRI in combination with T1-MRI, network-based statistics and a graph-theoretical approach to investigate differences between bilinguals and monolinguals in structural connectivity and topological properties of brain networks. Additionally, regional grey and white matter structural differences between groups were examined. The findings suggest that the effects of bilingualism on brain structure are not solid but unstable. However, lifetime experience of active bilingualism may lead to increased neural reserve in ageing, since better global network graph-efficiency has been observed in the elderly lifelong bilinguals compared to monolinguals.

## 1. INTRODUCTION

It has been proposed that bilinguals as compared to monolinguals show enhanced executive control (EC) functions (such as updating, switching, and inhibition) given the constant use of these mechanisms engaged in selecting, managing interference and inhibiting between languages^1^. Several behavioural studies supported this hypothesis by showing that bilingual perform better than monolinguals in EC tasks^2-4^. However, other behavioural studies reported similar performance for both bilinguals and monolinguals in the same tasks^5-11^. In recent years, neuroscientists have sought after neuroimaging evidence of brain changes to support this hypothesis. It is well known that experiences continuously change brain structure and functioning^12-14^. Therefore, brain differences could be expected when comparing bilinguals and monolinguals, especially in the language circuit. Nonetheless, if bilingualism leads to enhanced EC functions structural and functional differences should also expect in brain areas and circuits related to these EC processes.

The Adaptive Control Hypothesis (ACH) defines a general brain network for language control and bilingual speech production^15^. This network comprises the anterior cingulate cortex (ACC), pre-supplementary motor area (pre-SMA), the inferior frontal gyrus (IFG), the inferior parietal (including the supramarginal) gyrus, the basal ganglia (caudate and putamen), the thalamus, the insula, the cerebellum, and the premotor and motor cortex. The neuroimaging evidence shows structural brain differences between bilinguals and monolinguals^16-24^ consistently only in one of these regions: the IFG^19-21^. But the evidence also shows differences in regions not predicted by the ACH, such as the anterior inferior temporal gyrus^16,25,26^. Furthermore, some studies show opposite effects over the same grey matter (GM) regions (i.e., cerebellum^18,23,24^) and over the same white matter (WM) tracts (i.e., inferior frontal occipital fasciculus (IFOF) and corpus callosum^27-32^. Finally, other studies report not differences at all ^28,33,34^.

According to The ACH, different bilingual interactive contexts could produce a different kind of language switching behaviour and also change the target brain areas where to expect bilingualism effects. In short, bilinguals from dense code-switching language context do not need to employ EC mechanisms to deal with their languages as bilinguals from other dual-language environment do^15^. This fact would explain the lack of differences between bilinguals and monolinguals in some behavioural studies^5-7^. However, investigations with bilinguals from dual-language contexts without dense code-switching have also been unsuccessful at finding differences between both groups in tasks measuring EC functions^9-11^. Therefore, it is still not clear whether other factors explaining the absence of difference.

An additional factor that was brought into the debate is the sample age-selection. The majority of the studies on bilingualism have been carried out with young adults (20-40 years old). Having reached the top of development^41^, young adults would probably show a ceiling effect on some cognitive functions (such as EC) that would make differences between bilinguals and monolinguals more difficult to detect^42^. In fact, behavioural studies reported that bilinguals perform better than monolinguals in EC tasks in children, middle-aged adults and older adults, but less evident in young adults^43^.

Particularly, in the case of children, neuroimaging studies have been suggesting already that the neural responses pattern that supports language processing in infants is different in bilinguals than in monolinguals^44^ and that it involves different networks, particularly with greater connectivity to prefrontal areas in the bilingual group^45^. Considering also that the development of EC functioning occurs critically during early childhood^46^, extensive training in linguistic (and non-linguistic) skills could have a more profound impact on the brain that is more susceptible to changes due to the developing brain maturation^47^. Thus, if there are bilingual effects in the brain, they should be evident in children. So far, there are a reduced number of studies exploring variations in GM^48^ and WM^30,31^ structure among bilingual and monolingual children. Although these studies did not follow a whole-brain quantitative neuroimaging approach that could result appropriate to uncover differences in GM regions or WM tracts other than the targeted ones. Besides, changes in the anatomical connectivity of the brain related to bilingualism in childhood remain unknown, and variations in topological attributes of the brain network as well.

In the case of the older adults, the brain is declining and thus susceptible to neural compensation and neural reserve mechanisms^49^. Therefore, any potential difference should also be more likely to observe in the elderly population. Older adult bilinguals with more extended lifetime experience of active bilingualism might be more trained in language selection and thus in EC functions, which would boost brain regions related to EC^42^, becoming more resistant to atrophy and pathology. There is neuroimaging evidence suggesting bilingualism might produce a neuroprotective benefit in the brain against cognitive decline^16,17,28,29,50–52^. However, how bilingualism facilitates the neuroprotective effect (i.e., by neural reserve or compensation) is still poorly understood^53^. Here we explore these neural mechanisms by assessing the brain structural connectivity in lifelong bilinguals and monolinguals older adults. If lifelong bilingualism acts as a cognitive reserve variable, the configuration and the topological parameters of the brain network should reflect differences between elderly bilingual and monolingual groups. Bilingualism may develop specific subnetworks, which in turns help to compensate brain deterioration in old adult bilinguals (neural compensation) or may make the bilingual brain network (as a whole) more efficient in the use of their neural resources while becoming more resistant to brain deterioration (neural reserve).

Overall, there seems to be a strong case for investigating the effects of bilingualism on children and the elderly, as potential structural brain changes should be more clearly observed when cognitive functions are developing (in children) or declining (in elderly). Importantly, the bilingual population proposed for this study (Basque-Spanish bilinguals) is characterized as mostly simultaneous and early active bilingual that is continuously exposed to both languages and switches between these languages frequently in everyday life, and often interleave both languages in a single utterance within a conversation. This represents a dense code-switching language interactional context^15^. Moreover, both languages, Spanish and Basque, are very distant lexically^36^, orthographically^37^ and syntactically^38^. It is well known that age of acquisition (AoA) and language proficiency are determinant factors in the neural underpinnings of bilingualism^39^. Besides, when the properties of the two target languages differ more widely, the neural systems involved in the processing of both languages could be more segregated (i.e., may have distinct underlying neural correlates)^40^, which could also increase the possibility of finding structural brain changes in bilinguals. It is thus expected that the kind of bilingual population and the typological distance between languages, could increase the possibility of finding structural brain differences between groups.

As the neural mechanisms of bilingualism are instantiated by a large-distributed network^15,^ here we will use a large-scale neuroimaging approach. Due to the limitations intrinsic to each MRI technique, we will also combine two different Magnetic Resonance Imaging (MRI) techniques: T1-Weighted scans (T1-MRI) and Diffusion-Weighted MRI (DW-MRI) to investigate bilingualism related effects in the brain structure. DW-MRI will be used to determine the large-scale structural connectivity maps. For that purpose, the T1-MRI will be used to generate the GM parcellation employed in the connectivity analysis (e.g., 90 GM regions). For the data analysis, we will use a network-based statistic (NBS)^58^ and complex network analysis^59^ to isolate sets of regions interconnected differently and to identify differential topological properties of the networks, respectively. This methodology has been used before in the study of bilingualism in healthy young adults^61,62^. As in previous studies^61^, highly structural connected subnetworks in bilinguals as compared to monolinguals are expected: differently subnetworks obtained for the bilingual group will help capture the plastic changes in the language and EC networks. We also expect to find variations in the topological parameters (global/local efficiency) of the brain structural network and identify the IFG as a hub region in bilingual’s brain network, since this region appears consistently across both functional and structural studies. This network analysis represents a fine-grained spatial analysis of highly distributed systems in the brain.

Additionally, DW-MRI derived measures, such as fractional anisotropy (FA) will be used to identify regional WM differences using a tract-based spatial statistic (TBSS)^57^. The T1-MRI will be also used to reveal local GM volume differences between groups, using voxel-based morphometry (VBM) implemented in FSL software^54^. Due to differences observed in GM volume of cortical regions can be interpreted either as a difference in GM folding or thickness, a cortical thickness analysis will be performed using FreeSurfer^55,56^. Mainly, it is expected that bilingual show increased GM in brain regions that would be important for bilinguals living in a dense code-switching interactional language context (such as IFG, cerebellum, caudate/putamen). Regional increase WM is expected within the fiber bundles usually connecting these GM areas. This fusion between different neuroimaging techniques enables valuable complementary data in understanding structural brain plasticity, taking a more holistic and realistic approach to the whole brain to help clarify the neural bases of bilingualism.

## 2. RESULTS

### 2.1. GM analysis

#### 2.1.1.1 VBM

A mass univariate GLM was used, corresponding here to a two-way (2 × 2) between-factor ANCOVA adjusted for IQ as nuisance covariate, and we found a significant Language-Profile by Age-Group interaction effect at p < 0.005 TFCE corrected. One large cluster comprised an extended region including the right lingual, posterior cingulate (PC) and precuneus (see Table 3 and Figure 1). The post-hoc comparisons showed that the interaction effect in the right lingual/PC/precuneus was driven by the group of children. The children group showed a significant increase in GM volume for bilinguals compared to their monolingual peers (see also Table 3 and Figure 1).

**Figure 1.**
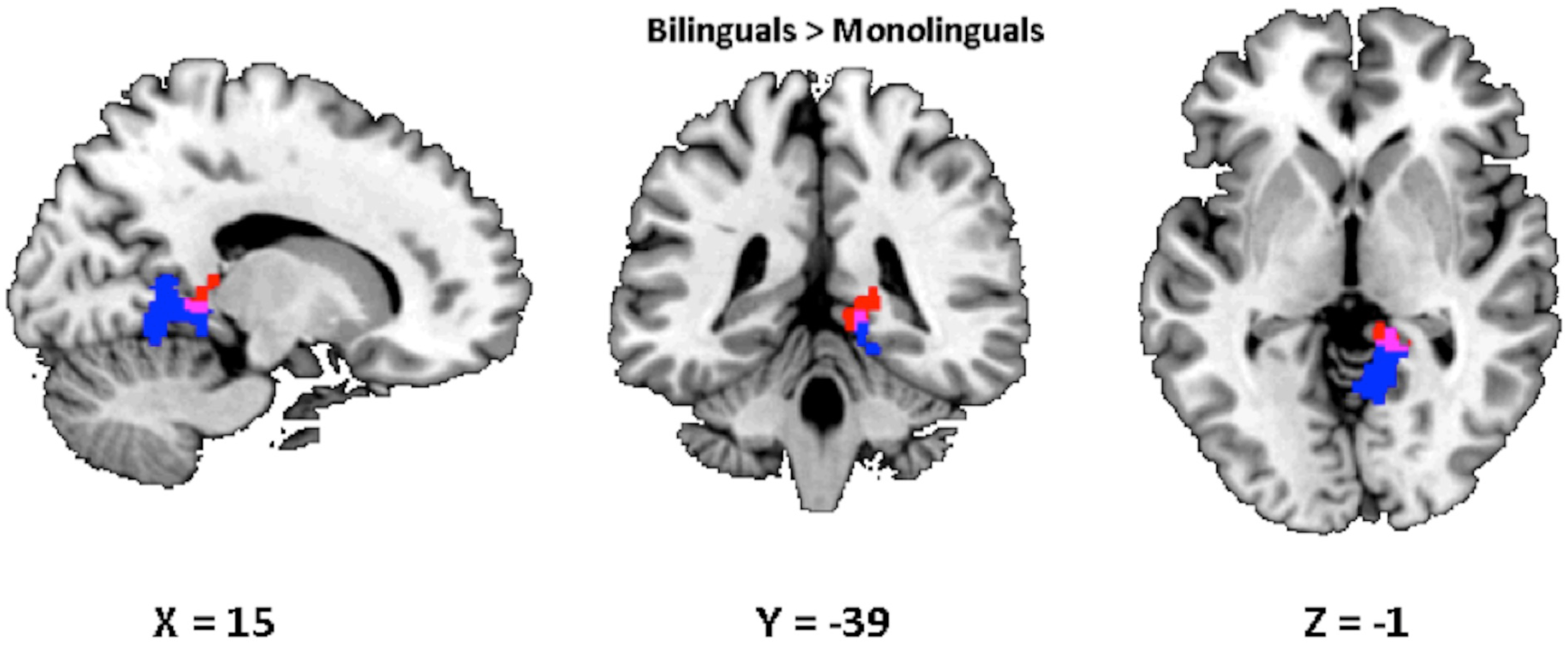
Brain regions in the right hemisphere showing significant increased GM in bilinguals compared to monolinguals. Significant cluster effects at p < 0.05 threshold-free cluster enhancement (TFCE) corrected. Red: Language-Profile by Age-Group interaction effect. Blue: Simple effect Bilinguals > Monolinguals for children. Purple: Overlay between the interaction and the simple effect. The background image is the MNI (Montreal Neurological Institute) brain template. Slices are taken from the X, Y, Z MNI standard coordinates (in mm) displaying the maximal overlay between the interaction and the simple effect, and are showing (from left to right): the sagittal, coronal and axial planes. The sagittal view represents the right hemisphere. In the coronal and axial views, the right hemisphere is on the right side. IQ as covariate, K = 10000 permutations. Bilinguals showed greater GM volume in the right precuneus, posterior cingulate and lingual gyri.

**Table 1.** Age in years, gender and IQ of children.

| **Monolingual group (N = 14, 6 female)** |  |  | **Bilingual group (N = 14, 6 female)** |  |  |
| --- | --- | --- | --- | --- | --- |
| **Age** | **Gender** | **IQ Composite (standard** | **Age** | **Gender** | **IQ Composite** |
| 6.36 | F | 141 | 6.42 | F | 103 |
| 8.32 | M | 100 | 8.54 | M | 109 |
| 8.30 | F | 102 | 8.35 | F | 102 |
| 9.20 | M | 121 | 8.73 | M | 114 |
| 10.57 | M | 110 | 10.45 | M | 106 |
| 10.98 | F | 100 | 10.71 | F | 108 |
| 11.05 | F | 124 | 10.74 | F | 106 |
| 12.10 | M | 101 | 11.86 | M | 105 |
| 12.78 | M | 100 | 12.79 | M | 104 |
| 13.10 | M | 115 | 12.95 | M | 102 |
| 13.16 | M | 122 | 13.35 | M | 111 |
| 13.25 | F | 116 | 13.51 | F | 109 |
| 13.61 | M | 119 | 13.78 | M | 90 |
| 14.14 | F | 117 | 14.22 | F | 109 |

*Note: N, sample size. IQ, intelligence quotient. F, female. M, male.*

**Table 2.** Age in years, gender and IQ of older adults.

| Bilinguals (N = 17, 10 female) |  |  |  | Monolinguals (N = 17, 10 females) |  |  |  |
| --- | --- | --- | --- | --- | --- | --- | --- |
| Age | Gender | IQ Composite<br>(standard scores) | MMSE | Age | Gender | IQ Composite<br>(standard scores) | MMSE |
| 66 | F | 123 | 29 | 66 | F | 123 | 29 |
| 69 | M | 119 | 29 | 69 | M | 117 | 27 |
| 68 | F | 107 | 30 | 68 | F | 120 | 28 |
| 76 | M | 131 | 30 | 76 | M | 87 | 28 |
| 70 | F | 117 | 29 | 69 | F | 102 | 29 |
| 75 | F | 90 | 28 | 75 | F | 85 | 28 |
| 67 | F | 118 | 30 | 67 | F | 87 | 29 |
| 78 | M | 98 | 27 | 78 | M | 99 | 30 |
| 71 | F | 85 | 30 | 71 | F | 86 | 30 |
| 73 | M | 103 | 29 | 73 | M | 110 | 30 |
| 70 | M | 125 | 26 | 69 | M | 110 | 29 |
| 68 | M | 92 | 29 | 68 | M | 84 | 28 |
| 65 | F | 123 | 30 | 65 | F | 95 | 28 |
| 64 | F | 122 | 30 | 64 | F | 89 | 30 |
| 69 | F | 120 | 28 | 69 | F | 113 | 26 |
| 66 | F | 125 | 28 | 66 | F | 83 | 28 |
| 65 | M | 133 | 29 | 65 | M | 105 | 30 |

*Note: N, sample size. F, female. M, male. IQ, intelligence quotient. MMSE, Mini-Mental State Examination scores.*

**Table 3.** Significant simple effect of Language-Profile in children showing increased GM volume in bilinguals as compared to monolinguals. The table is showing as well the Language-Profile by Age-Group interaction effect.

| Cluster Region | Voxels | P | X, Y, Z | T | *Interaction* |  |  |  |
| --- | --- | --- | --- | --- | --- | --- | --- | --- |
|  |  |  |  |  | *Voxels* | *P* | *X, Y, Z* | *T* |
|  |  | value | (mm) | value |  | value | (mm) | value |
| **R.Lingual/PC/Precuneus** | 1058 | 0.001 | 12, -58, -8 | 4.71 | 1235 | 0.007 | 2, -36, -8 | 4.9 |

*Note: PC, posterior cingulate; R, right; L, left; Voxels = number of voxels in the cluster; $P \leq 0.05$ threshold free cluster enhancement (TFCE) corrected. X, Y, Z = coordinates in Montreal Neurological Institute (MNI) space of local maxima. K = 10000 permutations. IQ as nuisance covariate.*

#### 2.1.2.1 Surface-based morphometry

In the vertex-based analysis of the CT, the ANCOVA showed two significant interaction between factors (Language-Profile by Age-Group) effects at p < 0.05 cluster-wise corrected using Bonferroni. These interactions were observed in the right precuneus extending into the lingual and PC (peak in MNI space [4.8 −58.1 22.1], p = 0.013, cluster size = 1298 vertices) and right postcentral gyrus (peak in MNI space [20.1 −32.2 59.1], p = 0.031, cluster size = 1307 vertices). The post-hoc comparisons showed that the interaction effects were driven by the children group, showing a significant thinner CT for bilinguals compared to monolinguals peers in the precuneus (peak in MNI space [8.4 −60.7 39.8], p = 0.006, cluster size = 3764 vertices). Also, a significant thinner CT in monolinguals compared to bilinguals in the postcentral (peak in MNI space [17.8 −32.5 59.0], p = 0.020, cluster size = 1440). See results in Figure 2.

**Figure 2.**
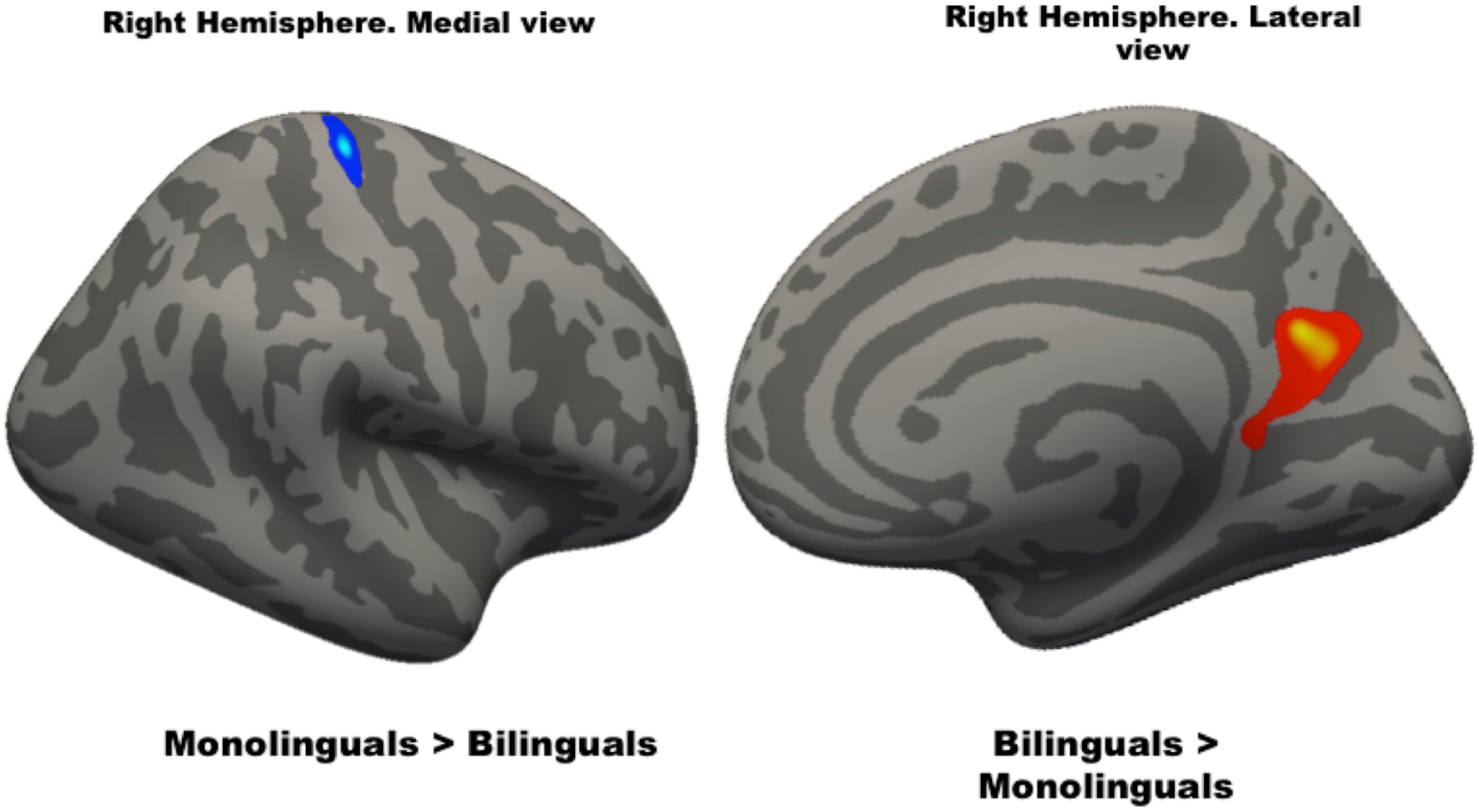
Brain region in the right hemisphere showing significant increased/decreased cortical thickness (CT) in bilinguals compared to monolinguals. ANCOVA results showing significant Language-Profile by Age-Group interaction effects at p< 0.05 Bonferroni corrected. Blue: Bilinguals > Monolinguals interaction driven by the group of children in the right postcentral. Red: Monolinguals > Bilinguals interaction driven by the group of children in the right precuneuos. The first and second background brain images are the lateral and medial view, respectively, of the right hemisphere inflated template. IQ as covariate, K = 10000 permutations. Right postcentral CT is increased in bilinguals relative to monolinguals and right precuneus CT is decreased in bilinguals relative to monolinguals. K = 10000 iterations.

#### 2.1.2.1.3 Relationship between GM effects and chronological age in children

Previous studies indicated that these regions (i.e. Lingual/Precuneus/CP) developed earlier during childhood^63^. Thus, in order to understand the increase GM volume and decrease CT in bilinguals as compared to monolinguals in the Lingual/Precuneus/CP cortices, we examined how the volume and CT in the cluster-regions of the effects were modified by the chronological age in a larger sample of 41 bilingual/monolingual children (20 females, age range, 6-14 years, mean age, 10.93 years, ±2.28 std). For this analysis the values of GM volume and CT inside the significant clusters of differences obtained from the whole-brain analysis were extracted an averaged. A Pearson correlation analysis between the age and the mean GM volume and CT extracted from the voxels inside these clusters was performed. No significant correlations were obtained across all participants between the structural measures and the age (r = 0.18, p = 0.27 for GM volume and r = 0.0001 p = 0.99 for CT).

We also examined if the two Linguistic-Profile groups: 25 bilinguals (13 females, age range, 6-14 years, mean age, 10.78 years, ±2.17 std) and 16 monolinguals (7 females, age range, 6-14 years, mean age, 11.18 years, ±2.50 std) show different age slopes in these cluster-regions. No significant correlations were obtained between these measures and the age for either bilinguals (r = 0.31, p = 0.13 GM volume and r = 0.068, p = 0.75 for CT) or monolinguals (r = 0.23, p = 0.39 for GM volume and r = −0.33 p = 0.20 for CT).

### 2.2. TBSS

The TBSS analysis of the FA exhibited a significant overall main effect of Language-Profile in the posterior part of the left IFOF and the inferior longitudinal fasciculus (ILF) at p < 0.05 TFCE corrected (see Table 4 and Figure 3). The FA values were overall decreased in these WM tracts for bilinguals.

**Figure 3.**
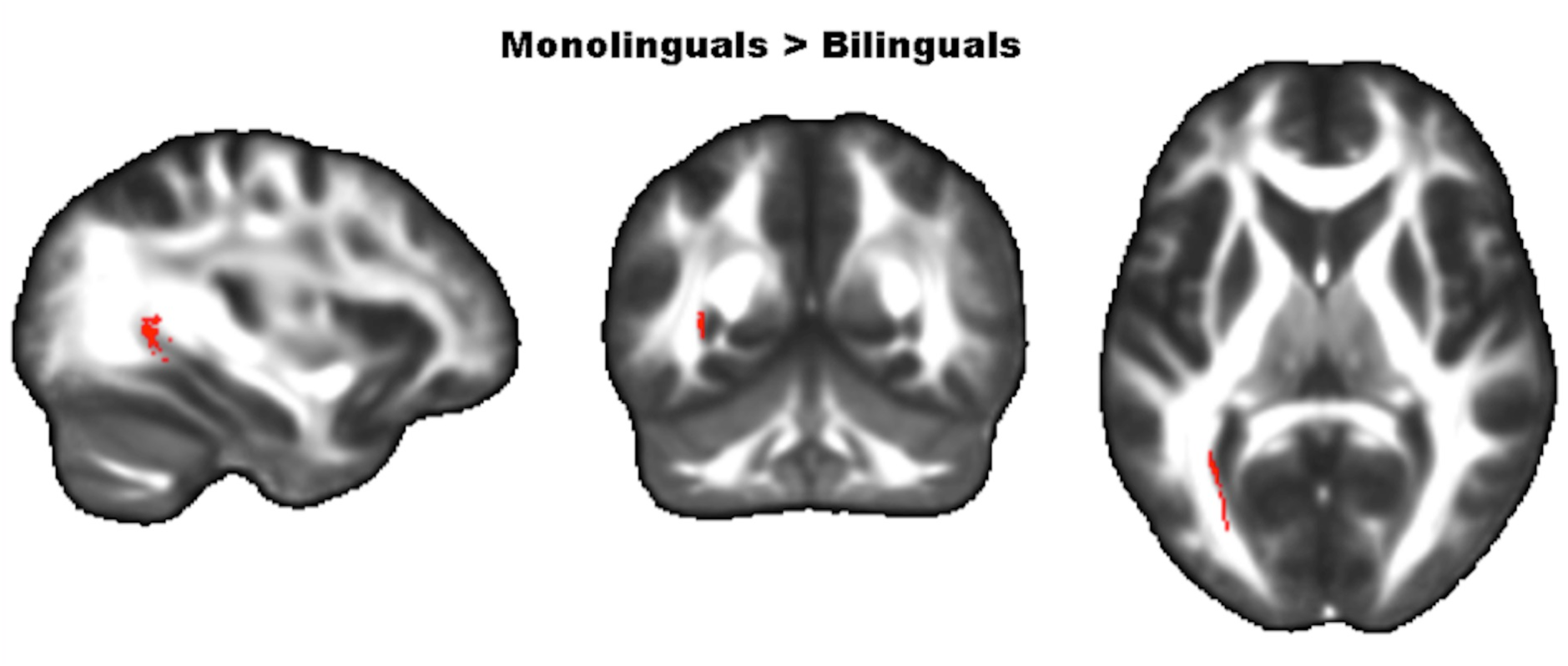
Brain regions showing significant increased fractional anisotropy (FA) in monolinguals compared to bilinguals in the left inferior frontal-occipital fascicule and inferior longitudinal fascicule. Significant cluster of overall main effect Monolinguals > bilinguals at p < 0.05 threshold-free cluster enhancement (TFCE) corrected. The background image is the FA brain template in MNI (Montreal Neurological Institute) space. The slices are showing (from left to right): the sagittal, coronal and axial plane. The sagittal view represents the left hemisphere. In the coronal and axial views, the left hemisphere is on the left side. IQ as covariate, K = 10000 permutations.

**Table 4.** TBSS analysis of the FA showing significant overall main effect of Language-Profile.

| Main Effect | Cluster<br>Region | Voxel | P-value | X (mm) | Y (mm) | Z (mm) | T-value |
| --- | --- | --- | --- | --- | --- | --- | --- |
| **Monolinguals > Bilinguals** | Left IFOF/ ILF | 268 | 0.039 | -32 | -53 | 3 | 4.32 |

*Note: $p \leq 0.05$ threshold free cluster enhancement (TFCE) corrected. X, Y, Z coordinates in Montreal Neurological Institute (MNI) space. K = 10000 permutations. IQ as covariate. L, left; IFOF, inferior frontal occipital fascicule; ILF, inferior longitudinal fascicule.*

### 2.3. Structural Connectivity

NBS (T-threshold = 3, K = 10000 permutations) did not identify any main effect of Language-Profile, or interaction between Language-Profile and Age-Group at p < 0.05.

### 2.4. Complex network analysis

The ANCOVA revealed a significant main effect of Language-Profile, F(1,56) = 6.29, p = 0.016 (see Figure 4) for E_glob_. Looking at the main effect could be noticed graphically that the lines are not parallel. That is because a significant interaction was also found between Language-Profile and Age-Group F(1,56) = 4.17, p = 0.047 for E_glob_. Post-hoc comparisons using Bonferroni correction indicated significantly higher E_glob_ for bilinguals compared to monolinguals in the elderly group but not in the children at p < 0.05 (see Table 5). This result suggested that the ability of the structural brain network to transfer parallel information between its regions was higher (or more efficient) in bilinguals than in monolingual. However, this efficiency only became significantly increased in the elderly bilinguals. No significant correlation was obtained between this measure and the MMSE scores (r = −0.025, p = 0.89). Finally, no significant effect at p < 0.05 were found for E_loc_.

**Figure 4.**
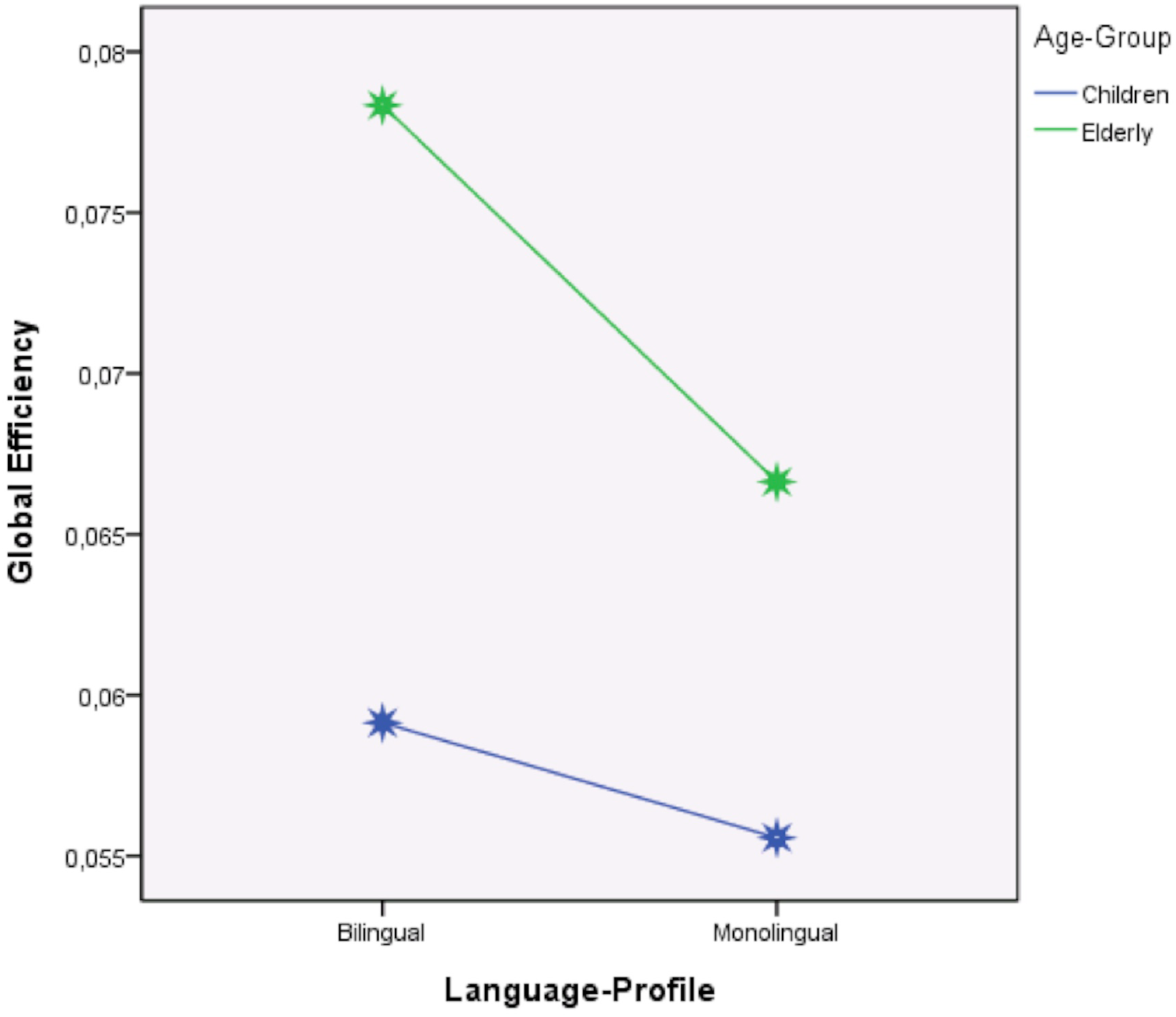
Graphical representation of the main effect of Language-Profile (x-axis) and the interaction between Language-Profile by Age-Group (separated lines, blue, children; green, elderly) on the global graph-efficiency (E_glob_) measure of the whole structural network (y-axis). Asterisks represent E_glob_ mean values for each group.

**Table 5.**
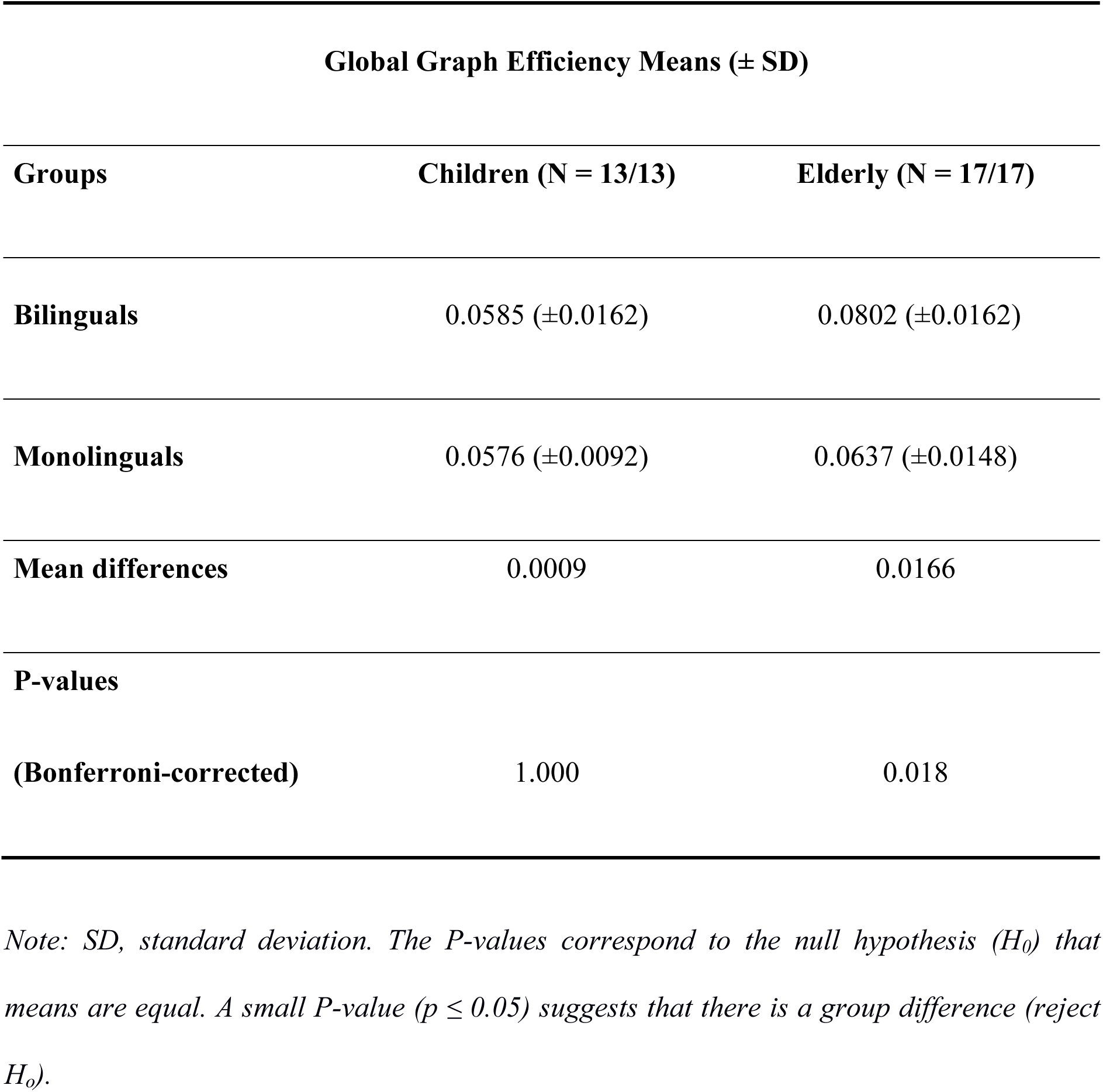
Global graph-efficiency comparison results between monolinguals and bilinguals.

Why is it the case that higher global network graph-efficiency is associated with overall reduced FA in the bilingual group? An increase in global graph-efficiency suggests that certain connections are stronger, which would lead us to expect greater density and higher FA values. However, this increased density may be masked by the existence of regions of crossing fibres. If a given tract strengthens its connections and becomes dense but intersects another tract that does not change, the FA values will be lower in the voxels of crossing fibres. This would explain why lower FA is related to increased E_glob_ of the brain graph-network. Nonetheless, these are measurements that rely on very different types of analysis. In fact, the higher global graph-efficiency could be related to more extensive changes that do not reach significance at a local level.

## 3. DISCUSSION

In the current study, different MRI measures such as GM volume, CT, FA, structural connectivity and topological parameters of brain networks (global/local graph-efficiencies) were combined to investigate brain plasticity changes in early active Basque-Spanish bilingual children and older adults. Both bilingual groups were carefully matched in age and gender with monolingual control groups. This study was carried out on samples of children and the elderly, with the aim of better understanding whether bilingualism yields plastic changes in the brain. Age groups were selected at either end of the lifespan under the assumption that any effects that bilingualism may produce in the brain would be more salient in these groups because they are not at the height of their cognitive skills (children are under development and elderly are on decline). Furthermore, it was expected that the potential differences obtained between bilinguals and monolinguals would be stronger in older adults than in children, given the impact of lifelong bilingualism in seniors compared to children. Predictions were made for differences in specific brain regions that might be important for bilinguals living in a dense code-switching interactional language context such as left IFG, caudate/putamen, cerebellum.

The study found a significant interaction between factors (Language-Profile by Age-Group) for the GM volume in the right lingual/PC/precuneus cortex and the CT in the right precuneus and postcentral gyri. The post-hoc comparisons revealed that the interactions were driven by the children group. They showed a significantly increased GM volume and a significantly decreased CT in the lingual/PC/precuneus for bilinguals compared to their monolingual peers. Additionally, bilinguals showed an increased CT in the postcentral gyrus compared to monolinguals. The TBSS analysis of the FA showed a significant overall main effect of Language-Profile in the IFOF/ILF. The FA was globally decreased in these WM tracts for bilinguals compared to monolingual peers. A significant main effect (bilingual > monolingual) for the E_glob_ of the structural network was also found, together with a less significant interaction between Language-Profile and Age-Group, suggesting that the whole brain structural network was more graph-efficient overall in bilinguals than in monolinguals, but mainly in the elderly group.

Contrary to what was expected, the elderly group did not show regional brain differences between Language-Profile groups. Although the recruited bilinguals were essentially simultaneous and early bilinguals, who have been immersed in a dense-code switching interactive bilingual environment for the majority of their life. Additionally, the extreme typological distance between the two languages (Basque and Spanish) might imply that the cost of dealing with them could have a greater impact in the brain, leading to more detectable plasticity changes in the bilingual brain. This result was in line with behavioural findings recently reported with the same elderly population^7^, showing no behavioural differences in EC tasks between bilinguals and monolinguals.

In contrast, children showed more regional brain changes than older adults, which could suggest that bilingualism might produce transient plastic changes in the brain when the languages are still being acquired, but later on, when they are established, these changes regress back to the previous state or diminish after the acquisition of the languages skills. This result is in line with the recent expansion and renormalization theory of brain plasticity in skill acquisition^64^. Specifically, bilingual children showed large increased GM volume and decreased CT in the right lingual/PC/precuneus. These patterns of results seem plausible since it has been previously demonstrated that the GM volume and the CT are inversely related^65^. These results are not compatible with the current neural model of bilingualism^15^ and with some prior structural findings^16-24^. However, the precuneus, as well as the PC, are essential regions in the ‘default mode network’^66,67^. These are crucial regions in development^68^ being the precuneus a region showing one of the highest indexes of maturation during childhood^63^ and one of the areas most connected in the brain^69,70^, although no correlation between the structural measures and the chronological age was obtained for children. However, evidence from prior studies indicates the involvement of the lingual, PC, and precuneus after oral language training in children with dyslexia^71^. Studies have also shown that Chinese-French bilinguals showed higher activations in the left cingulate gyrus and the right precuneus than the French monolinguals when performing a working-memory task^72^. And other study also found that a lexical decision task significantly correlated with the surface area of the left precuneus/PC gyri by studying a sample of highly proficient Spanish-Catalan bilingual^73^, which is consistent with the results demonstrating that these regions are associated with lexical and semantic processing^12,74^. Interestingly, one study in bilinguals have suggested the involvement of the precuneus and PC in language selection, set up, maintaining and updating the intention to speak in a particular language during a language-switching task^75^. The activations in precuneus/PC occurred before the lexical access or language execution phase (i.e. a preverbal stage). In general, our result suggests that bilingualism might influence these regions that are crucial for development and learning, especially for language selection, preparation and monitoring. Additional studies are required to explain the relationship of these neural correlates and bilingualism.

The study also showed an increased CT in the right postcentral gyrus for bilinguals, compared to monolingual children. There is a recent study^23^ found increased GM volume for Spanish-English bilingual young adults compared to English monolinguals in the right precentral gyrus covering part of the postcentral. However, they also found the inverse between simultaneous bimodal (ASL-English) bilinguals and monolinguals (i.e., decrease in the right precentral that extends into the postcentral for the bimodal bilinguals). Another study with Chinese learners of English^76^ found increased connectivity between the left postcentral gyrus and the right middle occipital gyrus in a pseudo-word rhyming task. These results suggest a potential role of the somatosensory cortex in learning languages.

Concerning to the analysis of the FA, the ANCOVA obtained an overall decreased FA for bilinguals across both groups of age: children and seniors. This decrease was located in the left IFOF/ILF. WM changes related to bilingualism have been observed consistently in the IFOF^27-32^. However, the results of previous WM studies are difficult to interpret. The effect may be pointing to an increase or a decrease in bilinguals. In any case, the overall main effect obtained in this study is consistent with result showing a reduction rather than an increase.

Contrary to what was expected, the NBS analysis of the structural connectivity did not show any set of regions with increased interconnectivity between Language-Profile groups. However, the topological analysis suggested that the whole structural brain network was more globally efficient in the elderly bilinguals than in elderly monolinguals, presumably due to the lifelong bilingual experience. The increased E_glob_ could be an indication that the early acquisition and the lifelong use of two languages could have a positive effect on the brain, allowing a more efficient flow of information across the whole brain network, which would, in turn, increase the brain’s ability to cope with focal deterioration in normal cognitive decline. Regardless, it is important to note that although these participants were likely in a declining process due to normal ageing, their cognitive states were at normal levels (Participants’ MMSE scores were above 26, mean = 28.76; std = 1.16) and did not differ between groups. Furthermore, the MMSE scores did not correlate with the E_glob_.

Notable, a previous study with young adults showed that early bilingualism developed higher interconnected and efficient subnetworks to deal with the processing of the two languages^61^. However, this change was associated with an observed decrease of the E_glob_ of the whole structural network. Those findings suggested that once the brain is more specialized and clustered (for example, forming dedicated subnetworks to manage with two languages) the E_glob_ tends to decrease. Conversely, the increased E_glob_ observed in the elderly bilinguals of the current study suggests that, perhaps, after the complete acquisition and extensive use of both languages in the elderly group, the brain specialization showed by young bilinguals could disappear and the brain becomes (or regresses back) more optimized, which might imply less specialization of the brain but better capability to transfer information across all brain network. Tentatively, this can suggest that lifelong bilingualism could act as a neural reserve mechanism, enabling the cognitive system to become more efficient at using cerebral resources and to deal better with the cognitive decline in ageing^64^. Nonetheless, the global graph network efficiency was the only significant effect observed in the elderly group and no relationship with participant cognitive state was observed. Thus, this finding needs to be replicated, and future investigations need to combine more behavioural measures with structural/functional measures to ascertain the relationship between this topological finding and his potential neural protective effect.

In conclusion, contrary to what was expected, bilinguals, as compared to monolinguals, showed more GM differences in children than seniors who have been active bilinguals during almost their whole lives. The effects of bilingualism on the structure of the brain in children were found in areas other than language and control regions, suggesting that bilingualism might accelerate the maturation in these regions (precuneus, PC and lingual gyri) during childhood that are crucial for development and learning. Although no relationship between structural measures and chronological age was observed. Finally, although lifelong bilinguals did not display regional differences in the structure of the brain, the brain could still show global differences. Lifelong bilingualism could point out gain toward an enhanced global graph-efficiency of the structural network in ageing, but this effect was not related with better participants’ cognitive state. In genreal, these results suggest that structural brain changes related to bilingualism could be unstable and difficult to detect even in lifelong bilinguals.

## 4. METHODS

### 4.1. Participants

#### 4.1.1 Children

Fourteen Spanish monolinguals (6 females, age range, 6-14 years, mean age, 10.98 years, 2.45 std) and 14 early Spanish–Basque bilinguals (6 females, age range, 6-14 years, mean age, 10.95 years, 2.48 std) participated in the study. The groups were paired in age and sex (see table 1). Participants were in good health and no reported history of neurological/mental illness or treatment with psychotropic medication. The parent of the participants gave verbal and written informed consent, following the Declaration of Helsinki, and the BCBL Ethics Committee approved the research protocol.

Bilingual children acquired both languages before preschool (mean of AoA, 0.23 years, ±0.83 std for Spanish and 0.91 years, ±1.5 std for Basque) and used both of them every day (daily exposure mean rates: 47.69, ±20.17 std for Spanish, 43.69, ±19.25 for Basque and 8.23, ±3.52 std for other languages). All of them born in the Basque Country, a region of Spain where both Basque and Spanish have co-official language status. We recruited bilingual children at the same bilingual school. In the Basque Country, bilingual schools ensure an equal amount of hours across languages and academic load. Children’s parents completed a language questionnaire and children were rated as highly proficient in both languages (mean rates: 9.57, ±0.53 std for Spanish, 8.20, ±1.39 std for Basque and 1.35, ±3.16 std for other languages) on a scale from 1 (the lower level) to 10 (the highly fluent). The scorings were based on reading, writing, listening, and speaking abilities. Monolinguals were recruited from other regions of Spain (Murcia, Cantabria and Asturias) where Spanish is the only official language. None of the monolinguals had fluent knowledge of other languages than Spanish (mean rate: 10, 0.0 std for Spanish) and did not belong to any immigrant minority. Monolingual and bilingual participants had a similar socioeconomic status. IQ scores were measured with the Spanish version of the Kaufman Brief Intelligence Test (K-BIT) and Wechsler Intelligence Scale for Children (WISC) and were controlled as a nuisance covariate. Notice that only 13 participants per group took part in the WM study because one participant from the bilingual group left the resonance before the acquisition of the DW-MRI sequence.

#### 4.1.2 Older adults

Thirty-four seniors who lived in the Basque Country were selected for this experiment (age range from 64-78, mean age, 69.35 years, ±4.01 std). They were recruited from Donostia–San Sebastián and had non-immigrant status. Participants were healthy people, with no reported history of neurological/mental illness or treatment with psychotropic medication and with normal or corrected-to-normal vision. All participants gave verbal/written informed consent, following the Declaration of Helsinki, and the BCBL Ethics Committee approved the research protocol.

The first group comprised 17 Basque-Spanish bilinguals (10 females, age range, 64-78 years, mean age, 69.41 years, ±4.08 std). The second group included 17 Spanish monolinguals (10 females, age range, 64-78 years, mean age, 69.29 years, ±4.07 std). They were paired in age and sex (see Table 2). The bilingual group rated themselves with a mid-to-high level of proficiency in both languages (mean rates: 8.83, ±1.02 std for Spanish and 7.70, ±1.48 std for Basque). The AoA of the second language (Spanish) ranged from 0 to 11 years old (mean age, 5.94 years, 2.92 std). All bilinguals had a lifelong bilingualism index (LB) above 91.48%, which means they have been bilingual for 91.84% of their lives (mean rates: 91.48, ±3.97 std for bilinguals and 4.58, ±18.90 std for monolinguals). This index represents an estimation of the amount of active exposure to both languages as a function of the age and is calculated from the formula: LBI = 100–(AoA L2*100/Age). In this way, both lately acquired bilingualism and short periods of active use of the two languages are represented by lower values, while early bilingualism and extensive use of the two languages get higher scores. Bilinguals used both languages every day. Monolinguals used only Spanish and had any/little knowledge of Basque (proficiency mean rates: 9.47, ±0.87 std for Spanish and 2, ±1.41 std for Basque) or any other language. Monolingual and bilingual participants had a similar socio-economic status.

The groups were not significantly different in mean years of study (mean rates: 18.88, ±5.78 std for bilinguals and 16.71, ±4.18 std for monolinguals) and cognitive state assessed by the Minimental State Examination Test (MMSE, Folstein 1975). MMSE mean scores: 28.88, ±1.17 std for bilinguals and 28.65, ±1.17 std for monolinguals. The study contained only right-handed old adults. IQ scores were measured with the Spanish version of the K-BIT and were controlled as nuisance covariate (mean rates: 113.58, ±14.84 std for bilinguals and 99.70, ±13.84 std for monolinguals).

### 4.2. MRI data acquisition

All images were acquired with a 3-T Magnetom Trio Tim scanner (Siemens AG, Erlangen, Germany). For each participant, 3D MPRAGE high-resolution T1-MRI scan was acquired. Acquisition parameters used were: echo time (TE) = 2.97 ms, repetition time (RT) = 2530 ms, 176 contiguous axial slices, voxel resolution 1 × 1 × 1 mm^3^, matrix size 256 × 256 and flip angle 7°. Total scan time was approximately 6 min.

DW-MRI data were recorded using a single-shot Stejskal-Tanner spin echo-planar-imaging (EPI) sequence with 64 gradient directions at b-value = 1500 s/ mm^2^ and 1 image at b = 0. The acquisition parameters were: TE/RT = 99/9300 ms, respectively, 58 contiguous axial slices, isotropic voxel resolution = 1.79 × 1.79 × 1.79 mm^3^, FOV = 230 mm, matrix size 128 × 128. Total scan-time was approximately 10 min.

### 4.3. Data pre-processing and analysis

In all the subsequent analyses, a 2 × 2 between-subject factors analysis of covariance (ANCOVA) was performed. The ANCOVA included two factors: Language-Profile (levels: bilinguals and monolinguals) and Age-Group (levels: children and elderly), adjusted for a covariate (IQ). Due to the high number of comparisons performed in the study the risk of false positive findings increased (Type I error). To minimize the Type I error, in addition the correction for multiple comparisons within each analysis, we performed FDR correction (q = 0.05) to set a critic p-value across analyses. All p-values less than or equal to the critic p of p = 0.047 were considered significant in this study^77^.

#### 4.3.1. Voxel-based morphometry

First, images were visually inspected for motion artifacts and were then automatic re-oriented to the avg152T1 template based on rigid-body realignment. Mainly, set the origin of the images (0 0 0 mm coordinates) to anterior commissure (AC) and correct head rotations. In this study, the FSL-VBM pipeline was used to analyze the structural data^53^ (http://fsl.fmrib.ox.ac.uk/fsl/fslwiki/FSLVBM), an optimized VBM protocol implement in FSL software^78^. The pre-processing of the structural images comprised brain-extraction, GM segmentation and non-linear registration^79^ into the MNI 152 standard space. The registered GM images were then averaged and flipped along the x-axis to create a symmetric group GM template. The GM segmented images in their native space were non-linearly transformed into the space of the group template, and modulated. Finally, the modulated registered GM segmentation images were smoothed with an isotropic Gaussian kernel with a sigma of 3 mm.

A voxel-wise general linear model (GLM) and permutation-based non-parametric testing^80^ was carried out (K = 10000 permutations). Regional differences were reported as significant at p < 0.05, corrected for multiple comparisons using threshold free-cluster enhancement (TFCE)^81^. An extent threshold of 50 voxels was also set. Anatomical labelling of significant regions was chosen from the MNI anatomical atlas^82^ integrated into FSL atlas tool and AAL atlas (Automated Anatomical Labeling)^83^ included in MRIcron software.

#### 4.3.2. Surface-based morphometry

Cortical thickness (CT) measures were obtained using FreeSurfer (version 5.1) (http://surfer.nmr.mgh.harvard.edu/). Cortical reconstruction includes: motion correction, skull tripped, automated transformation into Talairach space, subcortical WM and deep GM volumetric structures segmentation, tessellation of the GM and WM boundaries, automated topology correction and surface deformation that optimally place the GM/WM and GM/CSF borders^54,55^. Some deformations, including surface inflation and a high-dimensional nonlinear registration to a spherical atlas are applied. The segmentation and deformation algorithms produce representations of the CT that is the closest distance from the GM/WM boundary to the GM/CSF boundary at each vertex on the tessellated surface. The CT maps were smoothed using a 10 mm FWHM Gaussian filter.

The ANCOVA was performed using a vertex-wise GLM and non-parametric Monte Carlo testing (10000 iterations). A cluster-wise correction for multiple comparisons with initial cluster-formation threshold of t = 2 (p < 0.01) was used for statistical inferences. Clusters were considered as significant at p-value <0.05 corrected using Bonferroni.

#### 4.3.3. DW-MRI preprocessing

The DW-MRI dataset was pre-processed using FMRIB’s Diffusion Toolbox (FDT) implemented in FSL (available at http://www.fmrib.ox.ac.uk/fsl/). Using affine registration and the first volume in the dataset (b = 0 image) as a reference, eddy current distortions and head motion were corrected. Then, diffusion parameters were estimated in each voxel^78^. The FA images were obtained from this step.

#### 4.3.4. Tract-based spatial statistic (TBSS)

The analysis of the FA data was performed using TBSS^56^. The FA data were brain-extracted and aligned into the standard space using non-linear registration^79^. The mean FA was calculated and thinned to create a mean FA skeleton, and all individual registered FA were then projected onto this skeleton before the statistical analysis.

A voxel-wise GLM and permutation-based non-parametric testing (K = 10000 permutations) was performed. Differences were reported as significant at p < 0.05 corrected using TFCE with an extent threshold of 50 voxels. Anatomical locations of significant regions of effect were determined by the Johns Hopkins University (JHU) white-matter tractography atlas^84^ integrated into FSL atlas tool.

#### 4.3.5. White matter connectivity

For details in the connectivity analysis procedure see Garcia-Penton et al.^60^. Briefly, High-dimensional individual GM parcellations (90 GM regions) were generated from the T1-MRI to create the seed point masks used in the tractography. We estimated the connectivity values between each brain voxel and the surface of each of the seed-masks using the probabilistic fibre tractography implemented in FSL^85^. We used the voxel-region connectivity maps to create an individual undirected weighted connectivity matrix that represents the connectivity density between each pair of regions^86^. Then, we used the connectivity matrices to investigate differences in connectivity patterns between groups.

##### 4.3.5.1. Network-based statistic (NBS)

An NBS approach^57^ was used to perform a non-parametric statistical analysis on the large-scale structural brain networks created. The 2 × 2 between-subject factor ANCOVA was also used to identify the set(s) of interconnected regions (components/subnetworks) from the 90 × 90 connectivity matrices that differ significantly between the Linguistic-Profile groups. First, a two-sample T-test is performed independently at each edge of the connectivity matrix. We used a T-threshold = 3 to identify sets or components of supra-threshold edges. The number of edges that the components comprise is stored. Finally, a non-parametric permutation test (K = 10000 permutations) estimate the significance of each component^57,59^.

##### 4.3.5.2. Complex network analysis

We also investigated differences between groups in the global efficiency (E_glob_) of the whole-brain network and the local efficiency (E_loc_) of each node/region in the network. E_glob_ is defined as the inverse of the average shortest path length, where paths are sequences of linked nodes visited only once. So, decreased average shortest path length means a higher global efficiency of the network. The E_loc_ of a node is calculated as the E_glob_ in the vicinity of the node and shows how well its neighbours trade information when the node is removed. The E_loc_ is related to the grouping coefficient that represents to what extent a node is interconnected with the closest nodes^87^.

The same 2 × 2 between-subject ANCOVA was performed. For the E_glob_ one measure was obtained per participant, and for E_loc_ one measure was obtained for each 90 GM node/regions per participant. Significant effects were set at p < 0.05, corrected for multiple comparisons using Bonferroni.

## Code availability

All code can be requested from the corresponding author.

## Data availability

The datasets generated during and/or analyzed during the current study are available from the corresponding author on reasonable request.

## ACKNOWLEDGMENTS

We thank BCBL laboratory for their assistance with data collection; Margaret Gillon-Dowens, Perter Boddy, Karen Lopez, Brendan Costello, Natalia Drobotenko and David Soto for helpful edits and recommendations. This work was supported by grants PSI2015-65689-P, PSI2015-67353-R and SEV-2015-0490 from the Spanish Government, PI2015-1-27 from the Basque Government, AThEME-613465 from the European Union and an ERC-AdG-295362 grant from the European Research Council.

## AUTHOR CONTRIBUTIONS

LGP, JAD, MC conceived and designed the experiment. LGP, YF and AP preprocessed and analyzed the data. LGP, JAD and MC interpreted the results. LGP drafted the manuscript, which was reviewed, adapted and approved by all authors.

## CONFLICT OF INTEREST

The authors declare that there is no competing interests.

